# Active compensation for changes in *TDH3* expression mediated by direct regulators of *TDH3* in *Saccharomyces cerevisiae*

**DOI:** 10.1101/2023.01.13.523977

**Authors:** Pétra Vande Zande, Patricia J. Wittkopp

## Abstract

Genetic networks are surprisingly robust to perturbations caused by new mutations. This robustness is conferred in part by compensation for loss of a gene’s activity by genes with overlapping functions, such as paralogs. Compensation occurs passively when the normal activity of one paralog can compensate for the loss of the other, or actively when a change in one paralog’s expression, localization, or activity is required to compensate for loss of the other. The mechanisms of active compensation remain poorly understood in most cases. Here we investigate active compensation for the loss or reduction in expression of the *Saccharomyces cerevisiae* gene *TDH3* by its paralogs *TDH1* and *TDH2. TDH1* and *TDH2* are upregulated in a dose-dependent manner in response to reductions in *TDH3* by a mechanism requiring the shared transcriptional regulators Gcr1p and Rap1p. Other glycolytic genes regulated by Rap1p and Gcr1p show changes in expression similar to *TDH2*, suggesting that the active compensation by *TDH3* paralogs is part of a broader homeostatic response mediated by shared transcriptional regulators.

## Introduction

Biological systems are often robust to genetic and environmental perturbations (Félix and Barkoulas 2015; Gibson and Lacek 2020). This robustness is conferred in part by the presence of multiple genes in the genome with overlapping functions (Ohya et al. 2005; DeLuna et al. 2008; Diss et al. 2014). Such genes often arise through duplication events that give rise to two or more paralogous genes (Wagner 2000; Gu et al. 2003). As described in Diss et al. (2014), paralogous genes can contribute to phenotypic robustness through either passive or active mechanisms. In passive paralogous compensation, the normal activity of one of the paralogs is sufficient to minimize the phenotypic impact of losing the activity of the other paralog. By contrast, active paralogous compensation occurs when the activity of one paralog changes in response to loss of activity of the other paralog, reducing the phenotypic impact of this loss. For example, a gene may respond to loss of a paralogous gene’s function by increasing its expression level, producing more protein capable of performing the function of the mutated gene.

Multiple examples of active compensation by upregulation of a paralog have been identified (Rudnicki et al. 1992; DeLuna et al. 2010; Denby et al. 2012; Dong et al. 2016; Dohn and Cripps 2018; Rodriguez-Leal et al. 2019), but the molecular mechanisms responsible for such transcriptional compensation remain largely unknown. One notable exception is loss of the *CLV1* receptor kinase in *Arabidopsis thaliana* that is compensated for by the upregulation of related receptor kinases *BAM1, BAM2*, and *BAM3*. Under normal circumstances the *BAM* genes are negatively regulated by *CLV1*, and loss of *CLV1* removes this transcriptional repression, resulting in upregulation of the *BAM* genes that compensates for the loss of *CLV1* (Nimchuk et al. 2015). This active compensation for loss of *CLV1* is not conserved in tomato or maize, but other steps in the *CLV* signaling pathway show evidence of active or passive compensation within these species (Rodriguez-Leal et al. 2019). For example, in tomato, upregulation of *SlCLE9* in response to loss of *SlCLV3* reduces the phenotypic impact of the *SlCLV3* mutation, although the mechanism causing this upregulation is unclear (Rodriguez-Leal et al. 2019).

Large-scale synthetic genetic interaction studies in the baker’s yeast *Saccharomyces cerevisiae* have also shown that paralogs with overlapping function are frequently able to compensate for each other (Li et al. 2010; Kuzmin et al. 2020). Up-regulation of paralogous genes with overlapping functions when one paralog is deleted has been reported in *S. cerevisiae*, and paralogs with partially overlapping regulatory motifs are more likely to be dispensable than those without overlap suggesting compensation for their loss (Kafri et al. 2005). To explain these observations, a model has been proposed in which two paralogous enzymes that catalyze the same metabolic step and are coregulated by the same transcription factor form a network motif in which the accumulation of their metabolite, due to loss of one paralog, leads to upregulation of the other paralog, and thus active compensation (Kafri et al. 2005). There are many examples of feedback circuits from yeast to mammals with the potential to function this way, making the model potentially of wide relevance to many biological systems (Kafri et al. 2006). To the best of our knowledge, however, the proposed dependency on a shared regulator for active compensation by upregulation of paralogous genes has yet to be demonstrated empirically.

The *Saccharomyces cerevisiae TDH1, TDH2*, and *TDH3* genes are paralogs with overlapping protein function and partially overlapping regulation that might make them likely to show active compensation. All three of these proteins act as glyceraldehyde-3-phosphate dehydrogenases (GAPDHs) (McAlister and Holland 1985a; Linck et al. 2014), catalyzing a central step in both glycolysis and gluconeogenesis. The *TDH2* and *TDH3* proteins are most similar to each other, retaining 94% amino acid sequence identity (Holland and Holland 1980; Engel et al. 2014), whereas the *TDH1* and *TDH3* proteins have 89% amino acid sequence identity (Holland et al. 1983; Engel et al. 2014). *TDH2* and *TDH3* are expressed during exponential growth at different levels, and *TDH1* is expressed primarily during stationary phase (Delgado et al. 2001; Bradley et al. 2019). Deletion of *TDH3* reduces fitness to ∼90% of wild type whereas deletion of *TDH1* or *TDH2* alone has little to no effect (McAlister and Holland 1985b; Costanzo et al. 2010). The *tdh1М/tdh3М* double mutant shows a negative genetic interaction in which the double mutant is more deleterious than expected relative to the predicted additive effects of the two single mutant fitness measures (McAlister and Holland 1985b), and the *tdh2М/tdh3М* double mutant showed an even stronger negative interaction, growing at only 7% of wild type levels (Costanzo et al. 2010). These nonadditive impacts on fitness suggest that the functional overlap of these paralogs allows them to compensate for each other.

Here, we investigate the molecular mechanisms responsible for this compensation. We find that expression of both *TDH1* and *TDH2* are upregulated when *TDH3* expression is reduced, and downregulated when *TDH3* expression is increased, suggesting that both paralogs provide active compensation for changes in *TDH3* expression. However, *TDH2* was not upregulated in strains carrying mutations in direct regulators Rap1p or Gcr1p that decreased *TDH3* expression, suggesting that both Rap1p and Gcr1p are required for the compensatory upregulation of this paralog. For *TDH1*, compensatory changes in expression were seen in the Gcr1p but not Rap1p mutants, suggesting that there are differences in the molecular mechanisms providing compensatory changes in expression of the two paralogs. This involvement of Rap1p and/or Gcr1p in the upregulation of *TDH1* and *TDH2* provides empirical support for the model proposed by Kafri et al. (2005) in which active compensation by paralogous genes is facilitated by one or more shared regulators and feedback loops. This compensation is not limited to paralogous genes, however; we also see upregulation of other genes with shared regulators encoding proteins that function in the same metabolic pathway when mutations in the *TDH3* promoter reduce its expression but not in cells with reduced *TDH3* expression caused by mutations in Gcr1p or Rap1. Consequently, this study shows how shared regulators controlling expression of multiple (paralogous and non-paralogous) genes that function in the same biochemical pathway can provide mutational robustness through active compensation from other members of the pathway, contributing to homeostasis.

## Results

### Active compensation for loss of TDH3 by paralogs TDH1 and TDH2

To determine whether the compensation for loss of *TDH3* activity by *TDH1* and/or *TDH2* might be mediated by changes in their expression, we examined *TDH1* and *TDH2* expression in a *TDH3* deletion strain of *S. cerevisiae* (*tdh3Δ*) previously analyzed using RNA-seq (Vande Zande et al. 2022). We found that both genes showed significantly higher expression in the *tdh3Δ* strain than in the unmutated wild-type strain (Figure 1A, Wald test P-value for *TDH1* = 2×10^−5^, P-value for *TDH2* = 0.04). To determine whether the degree of upregulation correlates with the extent to which *TDH3* expression is altered, we used additional RNA-seq data from the same study to examine *TDH1* and *TDH2* expression in strains of *S. cerevisiae* carrying changes in the *TDH3* promoter that cause more moderate alterations in *TDH3* expression. Three of these strains carry a single point mutation in the *TDH3* promoter that drives either 20%, 50%, or 85% of wild-type *TDH3* expression (Vande Zande et al. 2022). A fourth strain carries a duplication of the *TDH3* gene with each copy carrying a single promoter mutation, resulting in a strain expressing *TDH3* at 135% of wild-type levels. We found that *TDH2* expression was negatively correlated with *TDH3* expression among these strains, with *TDH2* showing both increased expression when *TDH3* expression was decreased and reduced expression when *TDH3* expression was increased (Figure 1B). *TDH1*, on the other hand, showed more of a threshold-like relationship with *TDH3* expression: *TDH1* expression was strongly increased in the *TDH3* null strain, but only mildly (and similarly) increased in the mutant strains expressing *TDH3* at 20%, 50%, and 85% of wild-type levels (Figure 1C). Like *TDH2, TDH1* expression decreased in the strain overexpressing *TDH3* (Figure 1C). Taken together, these data provide evidence of active compensation when *TDH3* expression is altered, with expression of its paralogs *TDH1* and *TDH2* changing in ways expected to minimize the impacts of these *TDH3* mutations on fitness.

**Figure 1:**
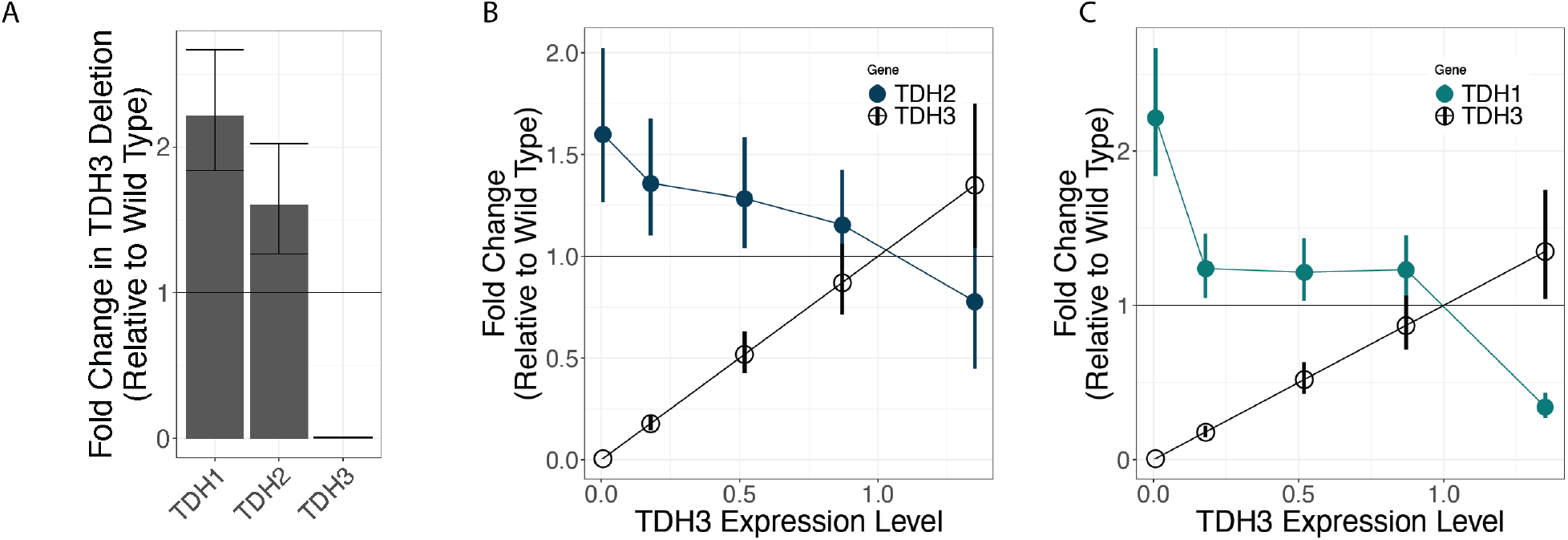
*TDH1* and *TDH2* actively compensate for changes in *TDH3* expression. (A) Changes in expression of *TDH1, TDH2*, and *TDH3* in response to the deletion of *TDH3* are shown, measured as fold change in expression relative to a wild type. Error bars represent one standard error of the mean. Statistical significance of expression changes was assessed using Wald tests in DESeq2, with the P-value for *TDH1* = 2×10^−5^, *TDH2* = 0.04, and *TDH3* = 7×10^−107^. Changes in expression of *TDH3* and *TDH2* are shown for strains with *cis*-acting mutations causing 0%, 20%, 50%, 85%, and 135% of wild type *TDH3* expression. Error bars show one standard error of the mean. (C) Changes in expression of *TDH3* and *TDH1* are shown for strains with *cis*-acting mutations causing 0%, 20%, 50%, 85%, and 135% of wild type *TDH3* expression. Error bars show one standard error of the mean.

### Active compensation might be caused by direct regulators of TDH3

For historical reasons (Duveau et al. 2017), the control strain and *TDH3* mutant strains profiled for expression using RNA-seq in Vande Zande et al (2022) all carried a reporter gene composed of the wild-type *TDH3* promoter allele driving expression of a yellow fluorescent protein (*P*_*TDH3*_*-YFP*). Surprisingly, we found that expression of this reporter gene was increased when native *TDH3* expression was decreased by mutations in its promoter and decreased by the duplication of *TDH3* with promoter mutations causing over-expression of *TDH3* (Figure 2A). This negative correlation between expression of the native *TDH3* gene harboring *cis*-acting mutations and expression driven by a wild-type allele of the *TDH3* promoter suggests that factors regulating expression of *TDH3* itself might be involved in the mechanism of active compensation.

**Figure 2:**
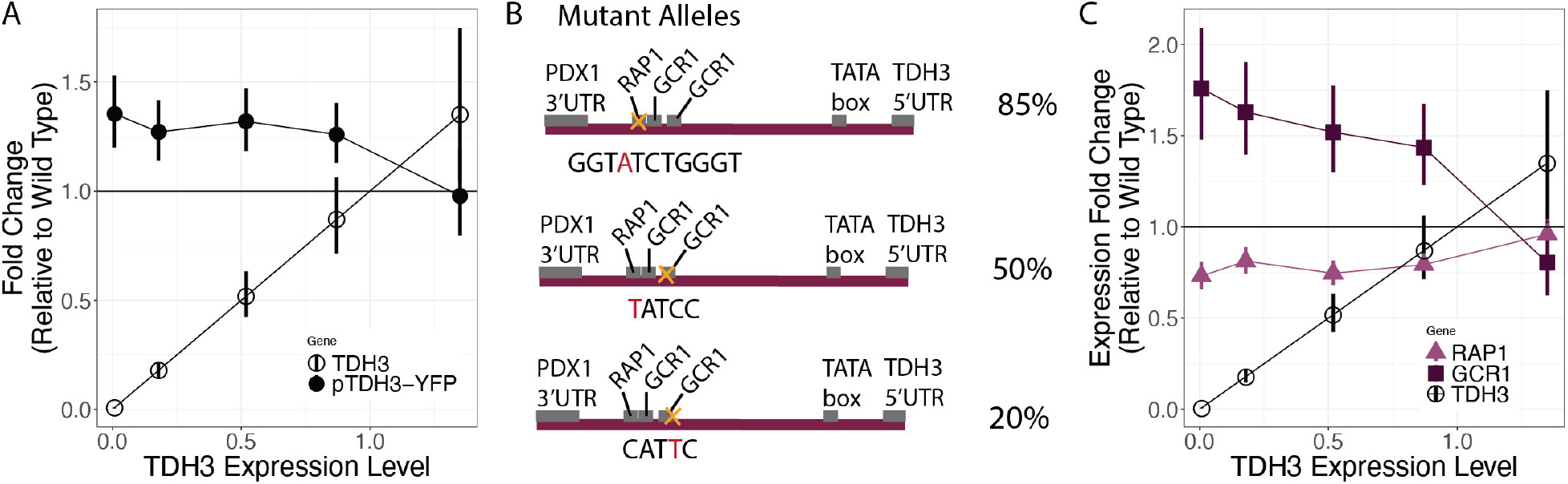
Feedback regulating TDH3 expression is mediated by RAP1 and GCR1 TFBSs. (A) Changes in expression of *TDH3* and a reporter gene with a wild type *TDH3* promoter driving expression of YFP (*P*_*TDH3*_*-YFP*) are shown for strains with *cis*-acting mutations causing 0%, 20%, 50%, 85%, and 135% of wild type *TDH3* expression. Error bars show one standard error of the mean. (B) Schematics and sequences of the *TDH3* promoter in mutant strains bearing mutations in binding sites for Rap1p and Gcr1p that result in *TDH3* expression levels of 20%, 50%, and 85% relative to wild type are shown. No schematic is shown for the mutant strain expressing *TDH3* expression at 135% of wild type levels, which contains two copies of the *TDH3* gene separated by a copy of the *URA3* gene, with both copies of *TDH3* containing a mutation in the binding site for Rap1p (GGTGTCTGaGT). (C) Changes in expression of *RAP1, GCR1*, and *TDH3* are shown for strains with *cis*-acting mutations causing 0%, 20%, 50%, 85%, and 135% of wild type *TDH3* expression, measured as fold change in expression relative to a wild type. Error bars represent one standard error of the mean.

The transcription factors Rap1p and Gcr1p regulate expression of *TDH3* (Huie et al. 1992; Yagi et al. 1994) as well as expression of other glycolytic genes, including *TDH1* and *TDH2* (MacIsaac et al. 2006; Hu et al. 2007; Venters et al. 2011; Lickwar et al. 2012). In fact, the mutations altering expression of *TDH3* in the mutant strains expressing *TDH3* at 20%, 50%, and 85% of wild-type expression levels all altered either Rap1p or Gcr1p binding sites in the *TDH3* promoter (Figure 2B, Duveau et al. 2017; Vande Zande et al. 2022). We thus wondered whether transcription of *RAP1* and/or *GCR1* was changed in the strains with *TDH3* promoter mutations. Using the same RNA-seq dataset described above, we found that *GCR1* was upregulated linearly in response to reductions in *TDH3* expression caused by mutations in the *TDH3* promoter whereas expression of *RAP1* was not (Figure 2C). If anything, expression of *RAP1* was slightly and similarly reduced in all mutants with reduced *TDH3* expression (Figure 2C). These observations suggest that changes in expression of *TDH1* and *TDH2* in response to changes in expression of *TDH3* might be caused by homeostatic feedback mechanisms involving direct regulators of *TDH3*.

*Mutations in RAP1 and GCR1 disrupt compensatory expression changes of TDH1 and TDH2* If Rap1p and/or Gcr1p are involved in the upregulation of *TDH1* and *TDH2* upon reduction of *TDH3* expression, we expect that strains with mutations in Rap1p or Gcr1p causing a reduction in *TDH3* expression would not show the same compensatory upregulation of *TDH1* and *TDH2*. That is, if the upregulation of *TDH3* paralogs requires Rap1p or Gcr1p, then mutations in these proteins that disrupt their ability to drive *TDH3* expression at wild-type levels should also impair their ability to upregulate expression of other genes in response to reduced *TDH3*. To test this hypothesis, we examined RNA-seq data from 9 mutant strains of *S. cerevisiae* each carrying 1-6 mutations in the *RAP1* (4 mutants) or *GCR1* (5 mutants) gene previously shown to affect *TDH3* expression (Duveau et al. 2021). These data were collected in parallel with the expression data for the *TDH3* mutants (Vande Zande et al. 2022). One *GCR1* mutant strain (GCR1.162) carried a single nucleotide deletion resulting in an early stop codon, suggesting it was likely to be a null mutation. This mutant expressed *TDH3* at only 7% of wild-type expression levels (Figure 3A).

**Figure 3:**
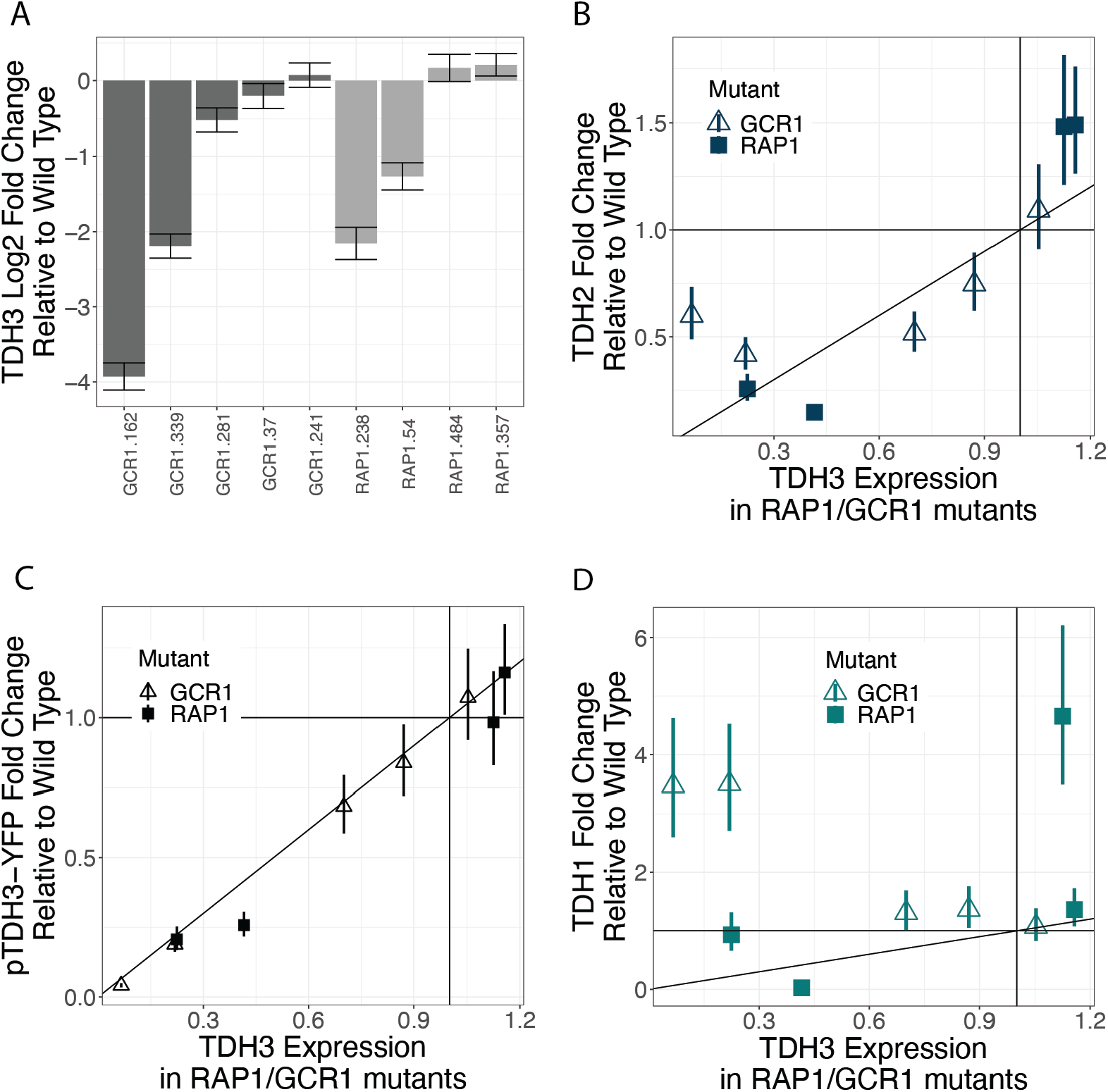
Mutations in *RAP1* and/or *GCR1* disrupt compensation by *TDH2* and *TDH1*. (A) Changes in expression of *TDH3* in response to various mutations in either *GCR1* (dark grey) or *RAP1* (light grey), measured as log_2_ fold change in expression relative to a wild type. Specific mutation identities in each strain are described in Table S1. Error bars represent one standard error of the mean. Fold changes in expression of *TDH3* and *TDH2* (B) a reporter gene with a wild type *TDH3* promoter driving expression of YFP (*PTDH3-YFP*) (C), and TDH1 (D), are shown for strains with mutations in either *RAP1* (squares) or *GCR1* (empty triangles). Error bars show one standard error of the mean.

The other *GCR1* mutant alleles were more likely to be hypo-(GCR1.339, GCR1.281, GCR1.37) or hypermorphs (GCR1.241), causing *TDH3* expression to range from ∼22% to ∼105% of wild type levels (Figure 3A). *RAP1* null mutants are lethal (Giaever et al. 2002), suggesting that all of the *RAP1* mutants examined were either hypo- or hypermorphs. These *RAP1* mutants showed *TDH3* expression ranging from ∼20% to ∼115% of wild-type levels (Figure 3A).

Consistent with Rap1p and Gcr1p mediating compensatory changes in paralog gene expression, we found that the *TDH2* gene was not upregulated in either the Rap1p or Gcr1p mutants that decreased *TDH3* expression (Figure 3B). *TDH2* expression was also not reduced in mutants causing overexpression of *TDH3* (Figure 3B). These observations indicate that both Rap1p and Gcr1p are required for the compensatory changes in *TDH2* expression seen in strains carrying mutations in the *TDH3* promoter. Changes in expression of the *P*_*TDH3*_*-YFP* reporter gene seen in the *TDH3* mutants (Figure 2A) were also absent in the *RAP1* and *GCR1* mutants altering *TDH3* expression (Figure 3C), again implying that Gcr1p and Rap1p were required for these changes. Expression of *TDH1*, on the other hand, showed compensatory increases in expression in *GCR1* mutants that lowered *TDH3* expression (Figure 2D), suggesting that Gcr1p is not required for the upregulation of *TDH1* in response to reduced expression of *TDH3*. Rap1p might be required for this active compensation, however, because neither of the *RAP1* mutants decreasing *TDH3* expression showed a compensatory upregulation of *TDH1* (Figure 2D). These data support a model in which Gcr1p is involved in the active compensation for changes in *TDH3* expression via *TDH2*, but not *TDH1*, with Rap1p involved in the compensatory changes in expression of both genes.

### Compensatory expression changes are also seen for other, non-paralogous, metabolic genes

Rap1p and Gcr1p are transcription factors that regulate expression of many metabolic genes (Uemura et al. 1997; Piña et al. 2003), thus active compensation for altered *TDH3* expression mediated by Rap1p and Gcr1p might affect more than just paralogous genes. Indeed, the eight genes encoding enzymes that function in the glycolytic pathway at steps immediately surrounding the step controlled by the TDH proteins have all been annotated as targets of Gcr1p and Rap1p based on either gene expression or chromatin immunoprecipitation experiments (Hu et al. 2007; Venters et al. 2011; Lickwar et al. 2012). We therefore examined the expression of these genes (Figure 4A) in the *TDH3, RAP1*, and *GCR1* mutants described above. We found that the genes *PFK2, PGK1*, and *ENO1* were significantly upregulated in the *thd3Δ* null mutant and their expression showed an inverse relationship with *TDH3* expression in the other TDH3 mutants examined (Figure 4B). Similar to *TDH2*, these compensatory changes in expression were absent when *TDH3* expression was altered by mutations in *RAP1* or *GCR1* rather than the *TDH3* promoter (Figure 4C). The genes *PFK1, ENO2, FBA1, TPI1*, and *GPM1* showed smaller changes in expression in the *TDH3* mutants and no statistically significant upregulation in the *thd311*. null mutant (Figure 4D). In the *RAP1* and *GCR1* mutants that altered *TDH3* expression, these genes showed expression similar to *TDH3* rather than compensatory changes in expression (Figure 4E). These expression patterns are consistent with the regulation of these glycolytic genes by Gcr1p/Rap1p as well as their active compensation for changes in *TDH3* expression being mediated through homeostatic feedback mechanisms involving Gcr1p and Rap1p.

**Figure 4:**
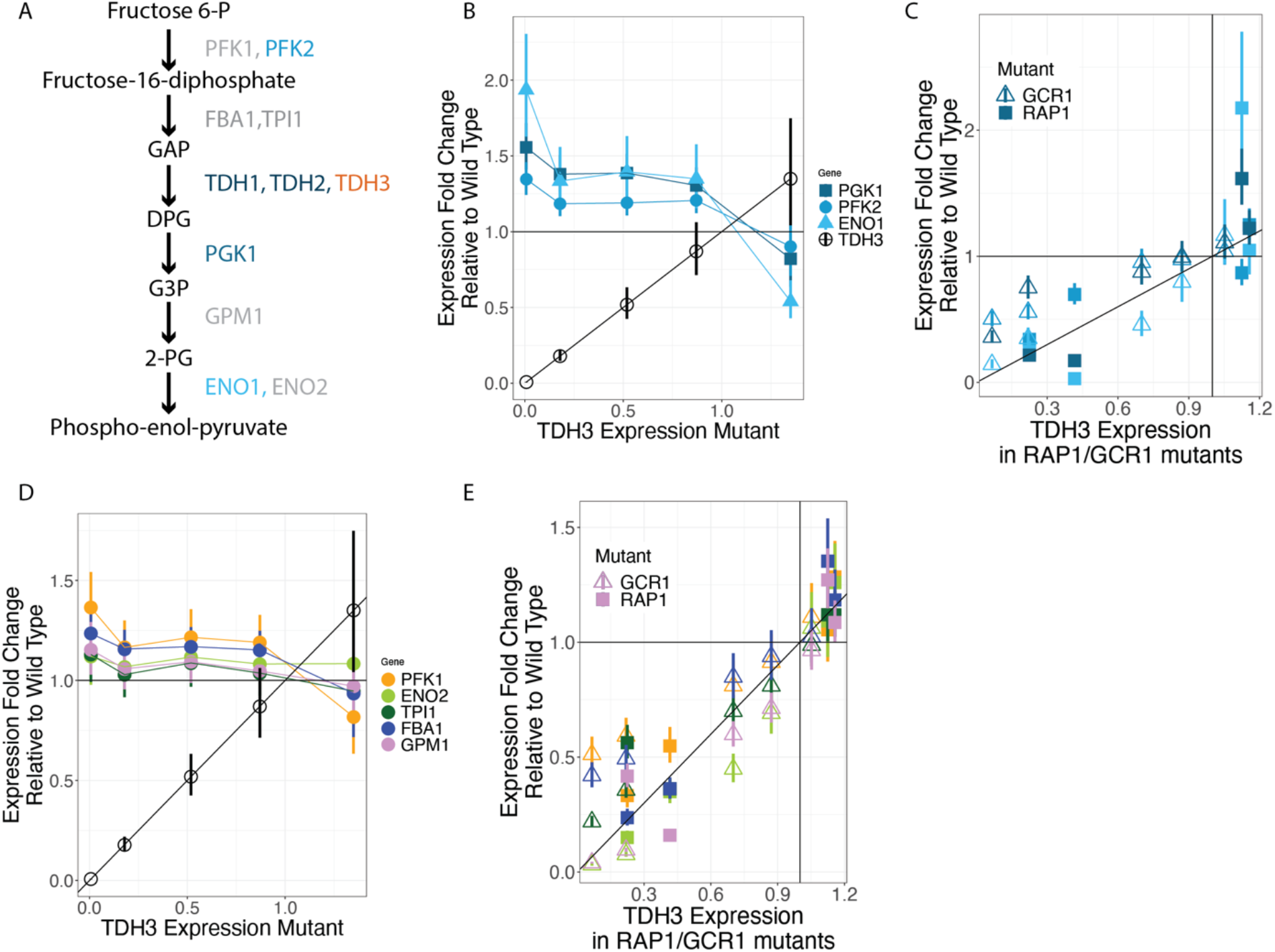
Multiple enzymes in the glycolysis pathway are upregulated upon reduction in *TDH3* expression in a RAP1/GCR1 dependent manner. (A) A simple schematic of the glycolytic pathway surrounding the metabolic step catalyzed by *TDH1,2*, and *3*, showing other enzymes catalyzing adjacent reactions. Enzymes that are significantly upregulated upon reduction in *TDH3* are in blue. Enzymes in this pathway that were not significantly upregulated are shown in grey. (B) Expression fold changes relative to wild type of the genes *PGK1, PFK2, ENO1*, and *TDH3* in yeast strains with varying levels of *TDH3* expression due to mutations in the native *TDH3* promoter, as estimated by RNA-sequencing data. Error bars are one standard error of the mean. (C) Expression fold changes relative to wild type of the genes *PGK1, PFK2, ENO1*, and *TDH3*, colored as in B, in the 9 yeast strains with varying levels of *TDH3* expression due to mutations in the genes encoding *RAP1* or *GCR1*, as estimated by RNA-sequencing data. Error bars are one standard error of the mean. (D) Expression fold changes relative to wild type of the genes *PFK1, ENO2, TPI1, FBA1, GPM1*, and *TDH3* in yeast strains with varying levels of *TDH3* expression due to mutations in the native *TDH3* promoter, as estimated by RNA-sequencing data. Error bars are one standard error of the mean. (E) Expression fold changes relative to wild type of the genes *PFK1, ENO2, TPI1, FBA1, GPM1* and *TDH3*, colored as in D, in 9 yeast strains with varying levels of *TDH3* expression due to mutations in the genes encoding *RAP1* or *GCR1*, as estimated by RNA-sequencing data. Error bars are one standard error of the mean.

## Discussion

Many genes with overlapping functions can compensate for each other’s loss, contributing to the genetic robustness of biological systems, but the mechanisms by which this compensation arises, operates, and is maintained over evolutionary time continues to be unclear (He and Zhang 2006; VanderSluis et al. 2010; Kuzmin et al. 2022). In this study, we show that changes in *TDH3* expression trigger feedback mechanisms that depend on the activity of transcription factors Rap1p and Gcr1p to offset the effects of these changes. Strains bearing *cis*-regulatory mutations in the *TDH3* promoter that decrease its expression presumably fail to upregulate *TDH3* because the transcription factor binding sites for Rap1p or Gcr1p are disrupted in these alleles (or because the locus is absent in the null mutant), yet expression of other genes regulated by Gcr1p and Rap1p is increased, including the *TDH3* paralogs *TDH2* and *TDH1* and even a reporter gene driven by a wild-type *TDH3* promoter. In other words, reduction in *TDH3* expression results in active compensation by upregulation of its paralogs, though seemingly through somewhat different mechanisms for *TDH1* and *TDH2* (Figure 5).

**Figure 5:**
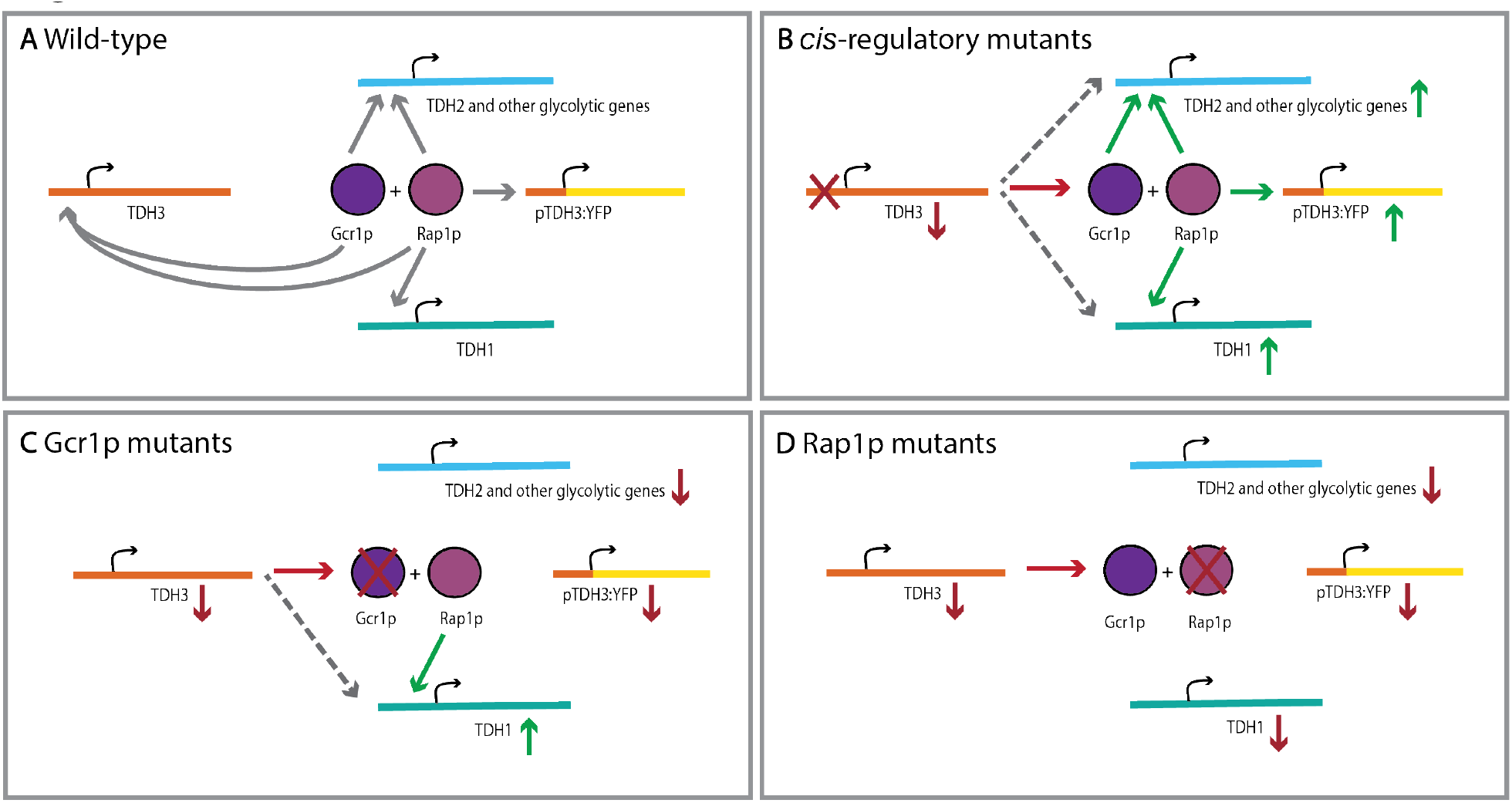
Model for active compensation by feedback and shared regulation. (A) In a wild-type cell, the Gcr1p and Rap1p complex regulate expression levels of *TDH2* and *3*, and Rap1p regulates expression of *TDH1*. (B) When the native promoter of *TDH3* is mutated, *TDH3* levels decrease, leading to an upregulation of *TDH2* and an intact *TDH3* promoter driving *YFP* via Gcr1p and Rap1p, and *TDH1* via Rap1p. (C) When Gcr1p is mutated, levels of all its direct targets are reduced. Lower levels of *TDH3* lead to an upregulation of *TDH1* via Rap1p. (D) When Rap1p is mutated, levels of all its direct targets are reduced. Despite lower levels of *TDH3* expression, the paralogs are not upregulated due to lack of functional Rap1p.

The upregulation of *TDH2* by Gcr1p/Rap1p might be achieved by increased expression of the *GCR1* gene in response to reduced *TDH3* expression. Transcriptional upregulation is not the only mechanism of activation of transcription factors (Hahn and Young 2011), but *GCR1* has been shown to be both transcriptionally and post-transcriptionally regulated by glucose availability (Hossain et al. 2016) and we observed increased *GCR1* expression in mutants with decreased *TDH3* expression, demonstrating that activity of this transcription factor is transcriptionally regulated under some circumstances. *RAP1*, on the other hand, performs roles in telomere maintenance and activation of ribosomal protein genes in addition to the activation of glycolytic genes (Sussel and Shore 1991; Shore 1994), and is not known to be transcriptionally responsive to metabolic changes. Because Rap1p and Gcr1p act in a complex to activate target gene expression, with Gcr1p being the major activator of the complex (Piña et al. 2003), we propose that upregulation of *GCR1* transcription upon reduction in *TDH3* expression is primarily responsible for the upregulation of the Rap1p/Gcr1p complex’s target genes, while still being dependent on functional Rap1p for upregulation of its target genes.

Active compensation by *TDH1* appears to occur via a different mechanism, as indicated by its more threshold-like response to reduction in *TDH3* expression and its upregulation in strains bearing mutations in *GCR1*. These differences in how *TDH1* and *TDH2* respond to reduction in *TDH3* expression may not be surprising since the expression pattern of *TDH1* has diverged from that of the other two paralogs (McAlister and Holland 1985a). *TDH1* has been shown to be upregulated under various stress conditions causing slow growth (Linck et al. 2014), and might therefore be upregulated by a mechanism related to the slower growth of mutants with reduced *TDH3* expression level rather than feedback specifically involving Gcr1p, although it does appear to be at least somewhat dependent on Rap1p function.

The fact that the upregulation of *TDH1* and *TDH2* does not completely eliminate the fitness effect of deleting *TDH3* suggests that pleiotropic effects of the compensation mechanism itself may carry a fitness cost (Kovács et al. 2021) and/or that the functions of these paralogs have diverged to some extent and cannot completely compensate for each other. Such partial subfunctionalization is thought to occur relatively frequently (Harrison et al. 2007; Kuzmin et al. 2020), and suggests that the maintenance of these paralogs by natural selection is not exclusively due to their ability to compensate for each other. Although *TDH3* is best known for its roles in glycolysis and gluconeogenesis, it has also been implicated in transcriptional silencing (Ringel et al. 2013), RNA-binding (Shen et al. 2014) and antimicrobial defense (Branco et al. 2014), functions which may not be able to be compensated for by *TDH1* or *TDH2* despite their high levels of protein conservation. More work assessing the ‘non-canonical’ or ‘moonlighting’ (Espinosa-Cantú et al. 2015; Chauhan et al. 2017; Singh and Bhalla 2020) functions of the GAPDHs in *S. cerevisiae* is needed to reveal the extent of subfunctionalization between these three paralogs.

The redundancy of paralogous genes, while imparting robustness to biological systems, simultaneously makes them unstable evolutionarily given that mutations in one gene are masked by the presence of the other gene. Yet, paralogous genes with overlapping function are maintained over long evolutionary timescales (Kafri et al. 2006; Tischler et al. 2006; Ihmels et al. 2007; Dean et al. 2008; DeLuna et al. 2008; Kafri et al. 2008; Hanada et al. 2009; Li et al.

2010; Kuzmin et al. 2020). Divergence in gene regulation and/or protein function might contribute to the maintenance of all three TDH paralogs over evolutionary time; however, in general, it remains to be seen how often the ability of paralogs to actively compensate for each other and contribute to genetic robustness is actively selected for or simply a side effect of their ancestrally shared regulators with sensitivity to feedback mechanisms. Decoding the molecular mechanisms responsible for active compensation among paralogous genes in other systems will help address this issue, revealing how living systems can thrive despite the inevitable changes in the environment and their genotype.

## Materials and Methods

### Strains used in this study

The *S. cerevisiae* strains used in this study are haploid strains derived from S288C and include the 5 *cis-*regulatory mutants affecting expression of *TDH3* containing changes in the *S. cerevisiae TDH3* promoter and the 9 *trans*-regulatory mutants affecting expression of *TDH3* that each carry 1-6 mutations in either the *RAP1* or *GCR1* gene described in Vande Zande et al. 2022. Construction of the *cis-*regulatory mutant strains, including the *tdh3*-***Δ*** strain, is described in (Duveau et al. 2017), and construction of the strains bearing mutations in the *RAP1* or *GCR1* genes is described in (Duveau et al. 2021). The collection numbers and specific mutations in each strain, as well as their impacts on *TDH3* expression, are detailed in Table S1.

### Gene expression data

RNA-sequencing data presented in this paper is a subset of the data described Vande Zande et al. 2022 and are available at GEO accession GSE175398. That dataset consists of RNA-sequencing data for *cis-*regulatory mutants and a larger set of *trans-*regulatory mutants affecting *TDH3* expression. Details of data collection and processing are available in (Vande Zande et al. 2022) and are summarized here. Briefly, yeast cells were grown to mid log phase in glucose media, pelleted, and frozen at -80C. polyA RNA was extracted from frozen cell pellets using oligodT magnetic beads. RNA libraries were prepared for sequencing using a ⅓ volume TruSeq RNA Sample Preparation v2 kit (Illumina) and sequenced on a HiSeq 4000 by the University of Michigan Sequencing Core. Each genotype (all mutants and non-mutated reference strains) was assayed in quadruplicate with each replicate consisting of a unique random array of genotypes and controls in a 96 well plate.

## Statistical analysis

All statistical analysis was performed in R, version 3.5.2). As described in Vande Zande et al. 2022, RNA-seq reads were pseudomapped to the *S*.*cerevisiae* transcriptome (Ensemble, release 38, retrieved from ftp://ftp.ensemblgenomes.org/pub/release-38/fungi/fasta/saccharomyces_cerevisiae/cdna/), and DeSeq2 (Love et al. 2014) was used to estimate log_2_ fold changes and significance values reported in the text. Code used in the analysis and to generate figures in this manuscript is available at Github (URL: https://github.com/pvz22/Compensation_TDH3).

## Acknowledgements

We thank Abigail Lamb for constructive feedback on the manuscript, Mo Siddiq and Holly Scheer for technical and intellectual support, and other members of the Wittkopp Lab for helpful discussions and feedback on drafts of this manuscript. This work was supported by the National Institutes of Health (grant number T32GM07544 to P.V.Z. and grant numbers R35GM118073 and R01GM108826 to P.J.W.) and the National Science Foundation (MCB-1929737 to P.J.W.).

